# Organ-on-Chip immunostaining method for three-dimensional identification and study of immune cells responding to drug-treated tumor cells

**DOI:** 10.1101/2024.04.24.590383

**Authors:** Francesco Noto, Adele De Ninno, Mario Falchi, Luca Businaro, Giovanna Schiavoni, Fabrizio Mattei

## Abstract

Epigenetic deregulation is implied in cancer initiation and resistance to antitumor drugs. In melanoma, aberrant DNA hypermethylation is frequently observed, resulting in the silencing of several genes involved in cell cycle regulation, apoptosis, tumor growth and drug resistance. DNA hypomethylating agents have been recently evaluated in both preclinical and clinical studies as a strategy to restore tumor suppressor genes and to increase immune recognition by tumors, highlighting their potential in pre-clinical models of melanoma. Advanced microfluidic system for the culture of complex three-dimensional cell, tissue and organ models have proven utility for oncoimmunology studies and drug testing. Here we present a protocol employing *ad hoc* fabricated microfluidic devices to reproduce a three-dimensional (3D) tumor microenvironment (TME) to study two aspects of the crosstalk between immune and cancerous cells under the effect of Decitabine (DAC), a DNMT inhibitor (DNMTi). First, we evaluated the preferential migration of immune cells towards treated and non-treated melanoma cells inside the chip. Next, we identified a specific subpopulation of migrated immune cells, with an on-chip immunostaining protocol resulting in the acquisition and evaluation of 3D images on a Laser-Scanning Confocal Microscopy (LSCM) station for in-depth characterization of tumor-immune interactions. This protocol may find broad application for pre-clinical drug testing in cancer studies.

## 1. Introduction

The benefit of cancer immunotherapies relies on the capacity of the host to develop a specific tumor immunity (Gracia-Hernandez et al., 2021; Marzagalli et al., 2019). Immune checkpoint inhibitors (ICB) have made a revolution in the treatment of cancer, but the relatively low percentage of patients responding to this therapy has raised the need for combinatorial approaches to improve its efficacy. Epigenetic modifier drugs display immune-related effects on tumor cells as well as on immune cells that could potentially synergize with ICBs. Epigenetic drugs such as DNMTi can improve antitumor activity depending on the effect on tumor immune landscape (Anichini et al., 2022). In melanoma, aberrant DNA hypermethylation is frequently observed, resulting in the silencing of several genes involved in cell cycle regulation, apoptosis, tumor growth and drug resistance. The epigenetic drug Decitabine (DAC), an inhibitor of the DNA methyl transferase enzymes (DNMTi), promotes immune recognition by re-activating silenced genes, in treating melanoma (Fazio et al., 2018; Giunta et al., 2021).

The recapitulation of organs and complex biological microenvironments by advanced 3D models, including organs on chip, have upgraded the conventional *in vitro* approaches with more accurate recapitulation of *in vivo* environments (Flont et al., 2023; Ma et al., 2021; Marconi et al., 2018; Mattei et al., 2021). In this regard, 3D organ on chip models have demonstrated potential for oncoimmunology studies, including the testing of drugs or drug combinations, and a valid tool to obtain parameters associated to the cross-talk between immune cells and cancerous cells (Businaro et al., 2013; Lucarini et al., 2017; Musella et al., 2022; Vacchelli et al., 2015).

We herein present an on-chip immunostaining protocol for the identification and 3D visualization of specific immune cell subsets interacting with cancer cells inside the chip chambers in a competitive cell migration assay. To this purpose, we employed a competitive on chip assay (Lucarini et al., 2017) to evaluate the preferential migration of peripheral blood mononuclear cells (PBMCs) towards A375M melanoma cells exposed or not to the DNMTi DAC. PKH67 green-labelled tumor cells are embedded in a Matrigel matrix, containing or not DAC, and loaded into separate lateral chambers while PKH26 red-labelled PBMC were loaded in a middle chamber. After 48 h of culture the preferential migration and infiltration of PBMC is towards either DAC treated vs untreated A375M gel containing chamber is evaluated by fluorescence microscopy. At this point, an immunostaning with anti-CD3 antibody (Ab) is carried out to identify CD3^+^ T cell subsets inside the two chip chambers (Matrigel-embedded melanoma cells with DAC or in absence of DNMTi drugs, respectively). After fixation of the microfluidic devices with a Paraformaldehyde/Glutaraldehyde mixture, the cells are ready to be acquired at high resolution, and 3D images (Z-stack) are captured by a LSCM station to perform spatial analysis of the CD3^+^ T cells on both chip chambers.

The innovation of this method relies on the potential to identify the phenotype of the specific immune cell subpopulations on chip chambers at the end of the dual TME competitive migration assay, as well as to analyze their precise 3D spatial distribution in the mimicked TME for more accurate evaluation of cancer-immune cell interactions.

## 2. Materials

### 2.1 Common Disposables

- 50 mL Polystyrene conical-bottom centrifuge tubes (#ET5050B, Euroclone)
- 15 mL Polystyrene conical-bottom centrifuge tubes (#ET5015B, Euroclone)
- 100 μm filters (Filcon sterile syringe-type (#340606, BD Biosciences)
- FACS tubes (e.g., 5 mL Polystyrene round-bottom tubes with cap, #352054, Corning)
- Sterile culture flasks with filter, 25 cm^2^ (#ET7026, Euroclone)
- Sterile culture flasks with filter, 75 cm^2^ (#ET7076, Euroclone)
- Cell count slides with counting grids (#87144, KOVA International)
- LS Columns (#130-042-401, Milenyi Biotec)

### 2.2 Cells and reagents

- Human melanoma cell line A375M (#CRL1619, ATCC)
- Buffy coats from human healthy donor peripheral blood (Prot. n. OO-ISS 02/10/2019 0029604 approval by the Ethics Committee of the Istituto Superiore di Sanità, Rome, Italy).
- DMEM High Glucose (#ECB7501L, Euroclone)
- Antibiotic-Antimycotic Solution (PSF) 100X (10000 U/mL Penicillin G, 10,000 μg/mL Streptomycin, 25 μg/mL Amphotericin B; #SV3007901, Euroclone)
- Fetal Bovine Serum (FBS) South America origin EU Approved (#ECS5000L, EuroClone)
- RPMI 1640 (#ECB9006L, EuroClone)
- Decitabine (DAC)(#A3656, Sigma-Aldrich, see *Note 1*)
- Phosphate-Buffered Saline (PBS)(#ECB4004L, Euroclone)
- Trypan Blue 0.4% solution (#17492E, Lonza, see *Note 2*)
- PKH26 Red Fluorescent Cell Linker (#MINI26, Sigma-Aldrich)
- PKH67 Green Fluorescent Cell Linker (#MINI67, Sigma-Aldrich)
- Biotin anti-human CD3 Antibody, primary antibody (#317320, BioLegend)
- Alexa Fluor 647 AffiniPure™ Goat Anti-Human IgG, F(ab’)_2_ fragment specific, secondary antibody (#AB_2337881, Jackson ImmunoResearch)
- DAPI (4’,6-Diamidino-2-Phenylindole, Dihydrochloride) (#D1306, Invitrogen)
- Matrigel Growth Factor Reduced Basement Membrane Matrix (#356231, Corning)
- Trypsin-EDTA in PBS (TEP)(#ECB3052, Euroclone)
- Bovine Serum Albumin (BSA)(#7030, Sigma-Aldrich)
- Paraformaldehyde (#441244, Sigma-Aldrich)
- Glutaraldehyde, 70% solution (#G7776, Sigma-Aldrich)
- Microfluidic Devices (see *Note 3*)
- Lymphosep, Lymphocyte Separation Medium (#AUL0560500, Aurogene)

### 2.3 Equipment

1. Micropipettes (0,5-10uL; 0-200 uL; 0-100uL) and polypropylene pipette tips
2. Class II Laboratory biosafety cabinet VBH 75 MP (#8071, Steril)
3. Forma Direct Heat CO_2_ Incubator (#310TS, Thermo Scientific)
4. LSCM station, LSM 900 model (#2651000312, Zeiss)(see *Note 4*)
5. SL 1R Plus MD bench centrifuge with full equipment (#17190589, Thermo Scientific)
6. Telaval 31 optical inverted microscope (#3514, Zeiss)
7. EVOS-FL Fluorescence microscopy station (#AMEFC4300, Invitrogen)

### 2.4 Software

1. Zen Blue v3.2 (Carl Zeiss GmbH)
2. ImageJ/Fiji v.1.54f with Java v1.8.0_172 (National Institutes of Health, USA; https://imagej.net/ij/)

## 3. Methods

### 3.1 Microfluidic device fabrication

We already set up a specific and standardized protocol for the fabrication of customized microfluidic devices, as already delineated in previous works (Mencattini et al., 2020). These microfluidic devices are made by using a specific biomaterial such as Poly-dimethylsiloxane (PDMS). It is a biocompatible silicone-derived elastomer, exhibiting excellent optical qualities and allowing direct observation by microscopy (Michielin & Maerkl, 2022; Wei et al., 2023; Yuan et al., 2023). Briefly, the microfabrication process can be resumed into i) fabrication of microstructured epoxy resin mold, ii) prototyping of PDMS channel microstructures by standard soft lithography, iii) bonding of PDMS chips to microscope slides or multiwell plates by oxygen plasma treatment and iv) chip functionalization poly-L-Lysine, if needed (see *Note 5*).

### 3.2 Tumor Cell culture

1. Human A375M cells are seeded in a 25 cm^2^ flask and routinely maintained in Complete Culture Medium (CCM, see Note 6) at 37°C and 5% of CO_2_.
2. The day of the experiment cells are detached from the flask with 1.2 mL of TEP solution, by incubation for 4 min at 37°C and 5% CO_2_ followed by a wash in 4 mL of Complete Culture Medium.
3. Cells are then counted and resuspended in a concentration of 1x10^6^ cells/mL. These cells will be used for PKH67 green cell tracker labeling.

### 3.3 Labeling of tumor cells with PKH67 Green fluorescent cell tracker

1. Follow the manufacturer’s instructions, with some minor changes. Wash A375M cells PBS 1X by centrifuging at 1500 rpm for 5 min at 4°C to eliminate every residue of supernatant. Discard the supernatant and resuspend cells in 500 μL of Diluent C. (see *Note 7)*.
2. *In an E*ppendorf vial resuspend 2 μL of PKH67 Red dye in 500 μL of Diluent C and rapidly add this mixture to the cell suspension. Incubate at r.t. (r.t.) for 5 min in the dark.
3. Stop the labeling process by adding 1 mL of FBS. Mix well and leave for 1 min at r.t.. Proceed to centrifuge at 1500 rpm for 10 min at 4°C.
4. Resuspend tumor cells in PBS 1X and count with Trypan Blue. Left PKH67^+^A375M green fluorescent cells at 4°C until use.

### 3.4 Isolation of PBMCs and labeling of tumor cells with PKH26 Red Fluorescent Cell Tracker

1. PBMCs are isolated from healthy donor’s buffy coats by centrifugation gradient method with Lymphosep, as previously reported in literature (Zırh et al., 2021).
2. At the end of the gradient separation procedure, resuspend the isolated PBMCs in PBS1X and count with Trypan Blue.
3. 12. To label the PBMCs with PKH26 Red Fluorescent Cell Tracker, the same A375M labeling protocol can be employed (see Section 3.3) as per manufacturer instructions included in the kit.

### 3.4 Microfluidic device Loading and acquisition

1. Prior of the loading, Matrigel stocks vials needs to be removed from -20°C freezer and placed in 4°C fridge, one day before the experiment (see *Note 8)*.
2. Before *s*tarting the experiment, place the Gilson’s pipette tips at 4°C fridge to maintain them cold during all the process (see *Note 8)*.
3. Prior to load cells (A375M and PBMC) into the chips prepare a mix with the Matrigel and the tumor cells for each condition. Namely, A375M cells to be exposed to DAC or to be left untreated are referred to as A375M-DAC. A375M cells or A375M-NT, respectively. Prepare two aliquots of Matrigel 2X, (concentration of 5 mg/mL), an aliquot of DAC 2X (concentration 0.5 μM) and a fourth vial containing CCM. Be sure that the final volumes used for these four aliquots are all the same (i.e., 100 μl).
4. Transfer the tumor cells in 1.5 mL vials and centrifuge at 1500 rpm for 5 min at 4°C (see *Note 9)*. The number of the cells to be used is based on their final concentration in the chamber. For an optimal culture, A375M are seeded at the concentration of 20000 cells in 5 μl Matrigel. After discarding the supernatant, cells are resuspended in the appropriate Matrigel mix depending on the experimental condition (NT or DAC). (see *Note 10)*.
5. Transfer tumor cells in a Eppendorf vial and centrifugate at 1500 rpm for 5 min 4°C. The number of the cells is based on the concentration of them in the chamber. For an optimal culture A375M are seeded in a number of 20.000 cells/chamber. After discarding the supernatant, cells are resuspended in the appropriate mix depending on the condition (see *Note 11)*.
6. After the A375-NT and A375-DAC samples are ready to be used, gently mix with a 10 uL pipette (see *Note 12) and load* in the inlet of the tumor chamber 5 uL of one solution (i.e., 5uL of the A375-NT mix; well 1 in Figure 1A and B) and 5 uL on the opposite tumor chamber of the other sample (i.e., 5 uL of the a375-DAC mix; well 2 in Figure 1A and 1B). To allow Matrigel propagation throughout the chambers 1 and 2 (Figure 1A), place the device upside down (with the plastic lid downwards) at r.t. for 5 min. At this point, transfer the chip (see *Note 13) in the in*cubator at 37°C and 5% of CO_2_ for 30 min (with the plastic lid upwards); this incubation does allow the gelation of Matrigel (see *Note 14)*.
7. In the meantime, PKH26^+^ PBMCs are prepared and resuspended in a concentration of 1x10^6^ cells in 10 uL of RPMI-CCM (see *Note 6)*.
8. After *s*olidification of gel matrix, hydrate all channels with 100 uL of CCM at r.t. (the medium is sequentially loaded in the wells 3, 4, 5, 6, 7 and 8; Figure 1A and 1B). At this point, the chips can be kept in the CO_2_ incubator until the loading of PBMCs starts (see Note 15). This will allow the formation of a 3D recapitulating TME (with or without drug) in the chip chambers delimited by the wells 1 and 2 (Figure 1A and 1B).
9. Lastly, to load the PBMCs in the central chamber, remove the medium from all the six wells (Figure 1A and 1B), and load 10 μL of cell suspension in the central channel (wells 3 and 4, Figure 1A and 1B). To refill the central chamber with the right amount of medium, 40 uL in the inlet were the PMBCs were loaded, and 50 uL in the opposite inlet.
10. Add 50 ul of CCM in the wells 5-6 and -8 (Figure 1A and 1B).
11. Using an EVOS-FL fluorescence microscope, we can check the preferential migration of the PBMCs into the side chambers. This does permit to evaluate the potential chemotactic activity of DAC compared to the competitive condition in the same chip (namely the A375M-NT chamber). To this purpose, acquisition microphotographs at 0 h, 24 h and 48 h time points will be done acquired by EVOS-FL microscope system to evaluate the presence of competitive migration of PBMCs (red fluorescence, Figure 1C).

**Figure 1.**
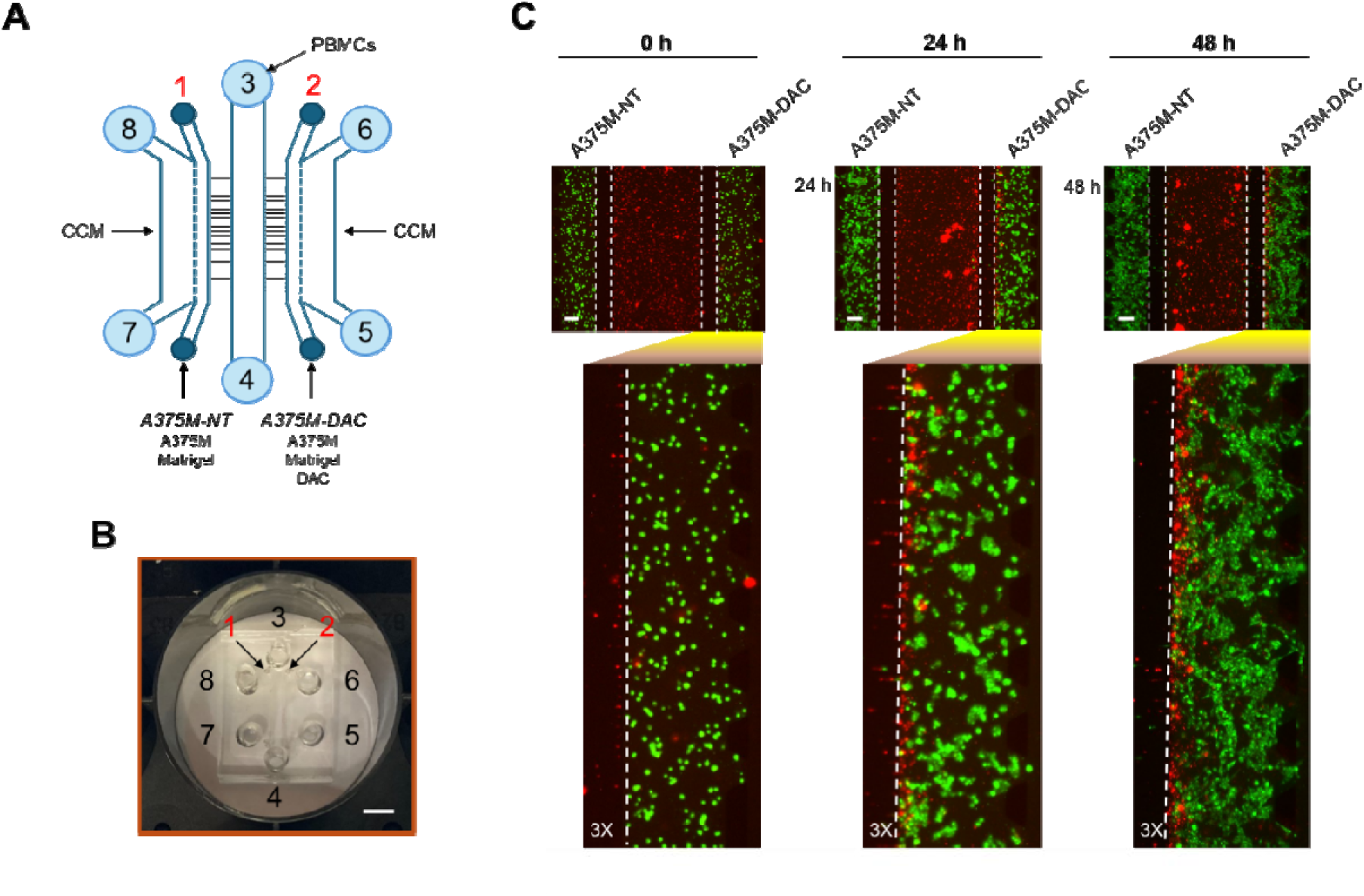
Overview of microfluidic chip loading pipeline and DAC activity on PBMC. (A) Loading workflow of tumor cells (wells 1-2; red color denotes that cells are resuspended in Matrigel) and PBMCs (wells 3-4) in a schematic representation of the microfluidic chip. Wells 5, 6, 7 and 8 are used to load CCM only and serves to balance the system. Text defines the experimental conditions and cells loaded in each well. (B) View of microfluidic device embedded on an optical support inside a standard 3x2 multiwell plate. Numbers delineate the wells employed for reagent and cell loading from panel A. Scalebar, 5mm. (C) Acquisition of microphotograph by EVOS-FL microscope at the indicated time points, with 3X magnification views (bottom panels). White dotted vertical lines delimit the microchannel arrays of the microfluidic device. Red fluorescence (PKH26), PBMCs. Green fluorescence (PKH67), A375M melanoma cells. Scalebar, 200 μm.

### 3.6 Labeling with anti-CD3 and secondary antibodies and fixation of microfluidic devices

1. Rinse the three upper wells 8, 4 and 6 (Figure 1A and 1B) with 50 μL PBS each, then repeat with the bottom wells (wells 7, 4 and 5, Figure 1A and 1B).
2. Discard the PBS from all the six wells (8, 3, 6, 7, 4 and 5, Figure 1A and 1B) and refill with a solution of PBS + 1% BSA. Leave the microfluidic device for 15 min at 4°C in the dark.
3. Remove the solution from all wells and proceed with the anti-CD3 Ab staining. Dilute the anti human CD3 Ab 1:200 in 300 μL PBS (see *Note 16) and load* sequentially the wells 8, 3, 6, 7, 4 and 5 (Figure 1A and 1B). Incubate for 45 min at 4°C in the dark.
4. Discard of the anti-CD3 Ab solution and wash with 50 μL PBS per well as previously described (step 1).
5. Stain with the secondary anti rabbit antibody Alexa Fluor 647 diluted 1:100 in 50 uL PBS per each well (total volume 300 μL). Load the six wells and leave for 45 min at 4°C in the dark.
6. Discard of the the secondary anti rabbit Alexa Fluor 647 solution and rinse with 50 μL PBS per well as previously described (step 1).
7. Load the fixating solution (PFA 2%, Glutaraldehyde 1% v/v in PBS) and incubate the chip 20 min in the dark at r.t.. (see *Note 17). Loading* sequence of the solution should start by wells 8, 3 and 6 and then wells 7, 4 and 5. Use 50 μL of fixating solution per well.
8. Aspirate the fixating solution and wash the chip in 50 uL of PBS 1X as described in the first point.
9. Proceed with the nuclei staining using the DAPI solution (diluted 1:250). Incubate the chip for 45 min at r.t. in the dark. Loading sequence of the solution should start by wells 8, 3 and 6 and then wells 7, 4 and 5. Use 50 μL of diluted DAPI solution per well.
10. Discard the DAPI solution and rinse in 50 μL of PBS 1X as described in the first point.
11. Add 50 μL PBS to each of the six wells. Do not add PBS to wells 1 and 2. At this point, the devices can be stored at 4°C up to 3 months.

### 3.7 Acquisition Z-stacks/XY planes microphotographs of microfluidic chip by LSCM platform

1. The Z-Stack/XY planes 3D microphotograps are acquired on a LSCM station Zeiss LSM 900 (Carl Zeiss GmbH, Jena, Germany) in Airyscan mode.
2. Excitation light was obtained by using diode lasers 405 nm, 488 nm, 647 nm and 555 nm. The optical thickness must be changed according to the objective used. It may range from 1 μm with a 10x objective till 0.2 μm with a 63x objective.
3. Images and Z-stack/XY planes (Figure 2) are obtained and processed by the Zen Blue (v3.2) software (Carl Zeiss GmbH, Jena Germany). Where needed, ImageJ is also employed for Z-stack/XY planes processing.

**Figure 2.**
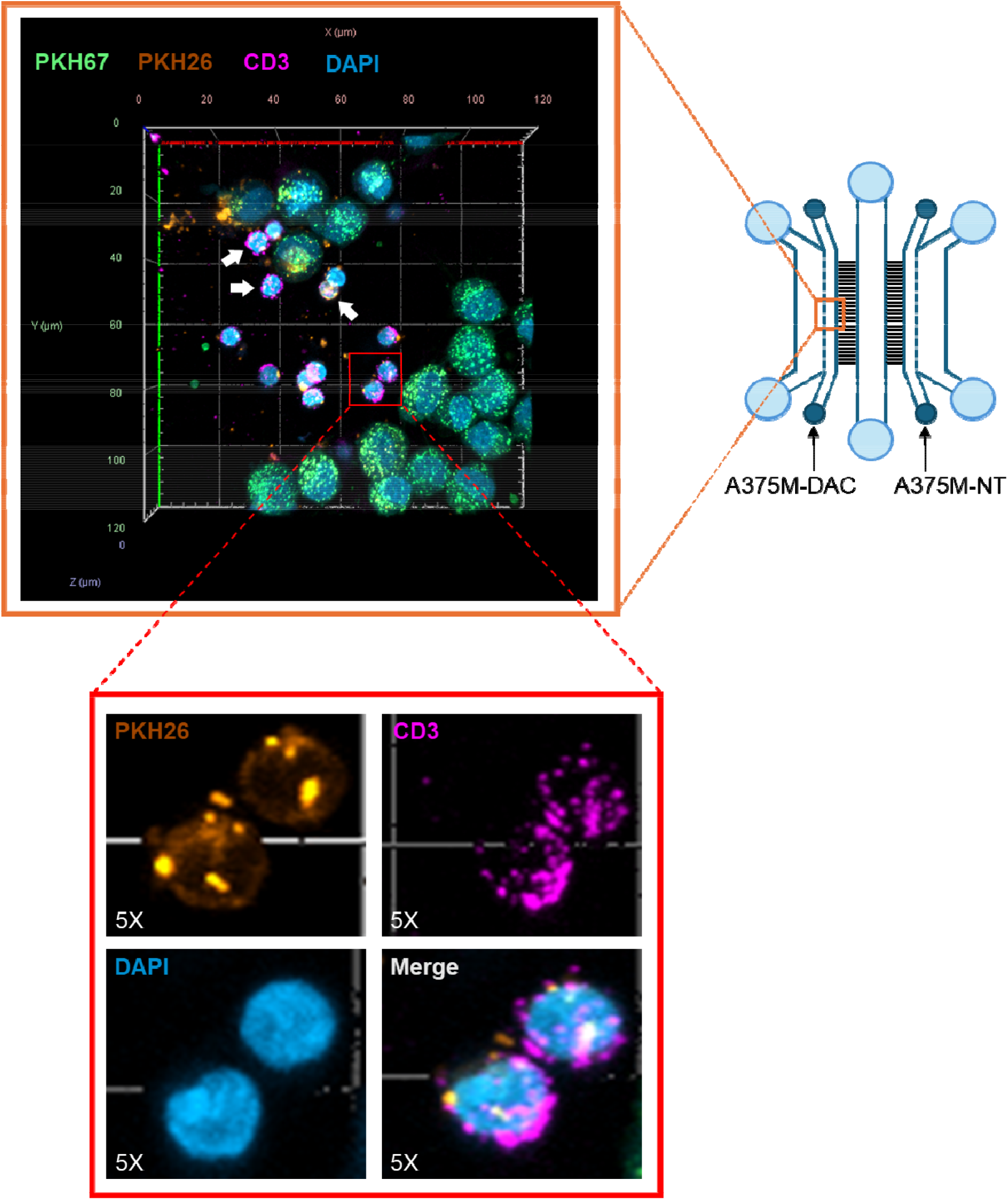
3D overview of T lymphocytes and melanoma cells from A375M-DAC chamber of the microfluidic chip at 48 h time point. Confocal analysis showing a Z-stack view on XY plane, acquired from DAC-treated A375M-gel chamber region (A375M-DAC, orange box), mimicking the DAC-shaped TME. This view details T lymphocytes (DAPI^+^PKH26^+^ CD3^+^ cells, with littler nuclei) spatially interacting (white arrows) or not with A375M melanoma cells (DAPI^+^PKH67^+^ cells, with bigger nuclei). Z-stack acquisition was performed by a 63X objective set up into the LSCM station. Bottom panel (red box) shows 5X magnification microphotographs with the fluorescence channels used to distinguish T lymphocytes from the resting PBMCs and melanoma cells.

## 4. Notes

1. We use DAC originally in a powder status. Ready-to-use DAC, is prepared by dissolving the powder in acetic acid and water with a stochiometric relation of 1:1 with the resulting concentration at 50mg/ml DAC. With this concentration stock we prepare 200 μL stock. Some aliquots of 100 uL volumes are stored at -80°C at this concentration and other equal volume aliquots are stored at a concentration of 10 mg/ml, corresponding to 44 mM DAC. Starting from the latter DAC dose, we achieved the final concentration of 0,25 μM, required for optimize the DAC epigenetic activity with a parallel minimization of its citotoxycity (Lucarini et al., 2017). The intermediate stock concentrations are needed to increase sensitivity during volume pipetting due to very little cell loading volumes for microfluidic chip. The other DAC stock can be used for more elevated working volumes.
2. To obtain a working solution dilute 1:8 in PBS and store at r.t..
3. Microfluidic devices are fabricated as previously specified in a clean room by lithographic methods (Businaro et al., 2013). The specific structures and substructures of the device used for these experiments is described in previous works (Lucarini et al., 2017; Musella et al., 2022).
4. The LSM 900 (3B class laser) LSCM station is equipped with Definite Focus 2.0 System, Axio Observer 7 Inverted Microscope (including a motorized Stage and optical objectives), CO_2_/O_2_ humidity/temperature controller (#11922, IBIDI), 2-Axes joystick CZ (#90762000820, Marzhauser-Wetzlar) and a Windows 10 operating system Desktop PC with the Zen Blue software.
5. Microfluidic devices are compatible with protein coating procedures (such as Poly-L-Lysine functionalization) depending on cell types and applications. If Matrigel is used as a gel matrix, there is no need to perform chip coating.
6. CCM is composed by DMEM High Glucose (500 mL), 1% Glutamine/Penicillin/Streptomycin/Fungizone mixture (PSF), 10% Fetal Bovine Serum (FBS). RPMI-CCM represents the RPMI-1640 medium enriched with the same additives of DMEM.
7. Diluent C is provided for both PKH26 and PKH67 labeling kits and must exclusively be employed to dilute the associated fluorochromes.
8. Matrigel is a gel matrix mixture that liquefy at low temperature (4°C) and starts to gelify at r.t. and over (for instance, 37°C). For these reasons, during the experimental procedures Matrigel must always be kept on ice. This assumption must be also be applied for all the material to be used to handle Matrigel.
9. Combine the two 2X Matrigel aliquots with the Complete Culture Medium and DAC 2X aliqutos, respectively. This does allow obtaining two a375M-NT (Matrigel plus A375M cells) and A375M-DAC (Matrigel plus A375M cells plus DAC) vials, with only the second aliquot containing the drug. Given that the initial four vials were prepared all with equal volumes, the final concentrations of Matrigel and DAC in these two vials will be 2.5 mg/mL and 0.25 μM, respectively.
10. Always plan loading of the microfluidic chips in at least duplicate per experimental condition. As a reference, a 100 μL solution is sufficient to load 20 side chambers of the chip for a total of 5 microfluidic devices. Consequently, the amount of solution to be prepared depends on the number of devices.
11. To obtain a 100 μL aliquot for the A375M-DAC, add to a 1.5 mL vial (containing 4x10^5^ pelleted A375M cells) 50 uL of 2X Matrigel (namely the 5 mg/mL Matrigel stock), 25 uL 2X DAC (namely the 0.5 uM stock) and 25 uL CCM. At the same time, to obtain a 100 uL vial for the A375M-NT condition add to 4x10^5^ pelleted cells 50 uL of 2X Matrigel and 50 uL Complete Culture Medium. These two 100 uL vial will be employed to load into the appropriate chip wells 20000 A375M cells per chamber.
12. Avoid to form bubbles during this mix process.
13. During the handling or transfer of the microfluidic devices, always manipulate them with extreme care.
14. This process involves the aggregation of Matrigel fiber components within the liquid phase to form a more ordinated network structure, resulting in a semisolid or gel-like substance. This represent a central step for the success of the microfluidic device loading process.
15. Maintaining the microfluidic device in the incubator at 37°C during unemployed times will ensure maintenance of Matrigel gel-like phase.
16. A volume of 50ul per each well of the device must be calculated. Therefore prepare 50 μL * 6 wells (corresponding to a total 300 μL 1:200 anti-CD3 Ab). The four little wells used to load Matrigel must not be considered. Indeed, solutions will spontaneously propagate in Matrigel chambers by passive gradient diffusion.
17. To obtain 500 μL of fixating solution mix 20 μL Glutaraldehyde (25%) to 250 μL of PFA (4%) and 230 μL of PBS 1X. Once used keep the solution at 4°C. The solution can be reused three or four other times.

## 5. Concluding remarks

Organ on chip and 3D models are innovating the study of solid tumor and their application has increased in biomedical research including oncoimmunology (Flont et al., 2023; Ma et al., 2021; Marconi et al., 2018; Mattei et al., 2021). The combined protocol herein described underlines the versatile nature of the organ-on-chip models. They represent an innovative tool of *in vitro recapitulat*ion of the TME, with a better accuracy compared the traditional 2D culturing techniques. The advantages of this Tumor-on-Chip approach (dual TME competitive migration assay) appear on the precise quantification of the preferential PBMCs infiltration in on chip generated competitive TMEs (DAC treated vs untreated) paralleled to the identification of a desired immune cell subset among the PBMCs. The identification and spatial localization of specific immune subsets within these reconstructed TMEs may allow, on the one hand, to unravel novel mechanisms of anti-tumor immune responses to treatments and, on the other hand, to tailor therapies according to the specific immune cell target.

## Acknowledgements

This work was supported by AIRC (Associazione Italiana Ricerca Cancro) IG 21366 to GS and in part by the Innovation Ecosystem Rome Technopole ECS00000024, funded by the European Union - Next Generation EU, PNRR Mission 4 Component 2 Investment 1.5.

## References

Anichini, A., Molla, A., Nicolini, G., Perotti, V. E., Sgambelluri, F., Covre, A., investigators, E. I.-o. C. A. E. (2022). Landscape of immune-related signatures induced by targeting of different epigenetic regulators in melanoma: implications for immunotherapy. J Exp Clin Cancer Res, 41(1), 325. 10.1186/s13046-022-02529-5

Businaro, L., De Ninno, A., Schiavoni, G., Lucarini, V., Ciasca, G., Gerardino, A., Mattei, F. (2013). Cross talk between cancer and immune cells: exploring complex dynamics in a microfluidic environment. Lab Chip, 13(2), 229–239. 10.1039/c2lc40887b

Fazio, C., Covre, A., Cutaia, O., Lofiego, M. F., Tunici, P., Chiarucci, C., Maio, M. (2018). Immunomodulatory Properties of DNA Hypomethylating Agents: Selecting the Optimal Epigenetic Partner for Cancer Immunotherapy. Front Pharmacol, 9, 1443. 10.3389/fphar.2018.01443

Flont, M., Dybko, A., & Jastrzębska, E. (2023). A layered cancer-on-a-chip system for anticancer drug screening and disease modeling. Analyst, 148(21), 5486–5495. 10.1039/d3an00959a

Giunta, E. F., Arrichiello, G., Curvietto, M., Pappalardo, A., Bosso, D., Rosanova, M., On Behalf Of Scito Youth. (2021). Epigenetic Regulation in Melanoma: Facts and Hopes. Cells, 10(8). 10.3390/cells10082048

Gracia-Hernandez, M., Munoz, Z., & Villagra, A. (2021). Enhancing Therapeutic Approaches for Melanoma Patients Targeting Epigenetic Modifiers. Cancers (Basel), 13(24). 10.3390/cancers13246180

Lucarini, V., Buccione, C., Ziccheddu, G., Peschiaroli, F., Sestili, P., Puglisi, R., Mattei, F. (2017). Combining Type I Interferons and 5-Aza-2’-Deoxycitidine to Improve Anti-Tumor Response against Melanoma. J Invest Dermatol, 137(1), 159–169. 10.1016/j.jid.2016.08.024

Ma, C., Peng, Y., Li, H., & Chen, W. (2021). Organ-on-a-Chip: A New Paradigm for Drug Development. Trends Pharmacol Sci, 42(2), 119–133. 10.1016/j.tips.2020.11.009

Marconi, A., Quadri, M., Saltari, A., & Pincelli, C. (2018). Progress in melanoma modelling in vitro. Exp Dermatol, 27(5), 578–586. 10.1111/exd.13670

Marzagalli, M., Ebelt, N. D., & Manuel, E. R. (2019). Unraveling the crosstalk between melanoma and immune cells in the tumor microenvironment. Semin Cancer Biol, 59, 236–250. 10.1016/j.semcancer.2019.08.002

Mattei, F., Andreone, S., Mencattini, A., De Ninno, A., Businaro, L., Martinelli, E., & Schiavoni, G. (2021). Oncoimmunology Meets Organs-on-Chip. Front Mol Biosci, 8, 627454. 10.3389/fmolb.2021.627454

Mencattini, A., De Ninno, A., Mancini, J., Businaro, L., Martinelli, E., Schiavoni, G., & Mattei, F. (2020). High-throughput analysis of cell-cell crosstalk in ad hoc designed microfluidic chips for oncoimmunology applications. Methods Enzymol, 632, 479–502. 10.1016/bs.mie.2019.06.012

Michielin, G., & Maerkl, S. J. (2022). Direct encapsulation of biomolecules in semi-permeable microcapsules produced with double-emulsions. Sci Rep, 12(1), 21391. 10.1038/s41598-022-25895-8

Musella, M., Guarracino, A., Manduca, N., Galassi, C., Ruggiero, E., Potenza, A., Sistigu, A. (2022). Type I IFNs promote cancer cell stemness by triggering the epigenetic regulator KDM1B. Nat Immunol, 23(9), 1379–1392. 10.1038/s41590-022-01290-3

Vacchelli, E., Ma, Y., Baracco, E. E., Sistigu, A., Enot, D. P., Pietrocola, F., Kroemer, G. (2015). Chemotherapy-induced antitumor immunity requires formyl peptide receptor 1. Science, 350(6263), 972–978. 10.1126/science.aad0779

Wei, Y., Wang, T., Wang, Y., Zeng, S., Ho, Y. P., & Ho, H. P. (2023). Rapid Prototyping of Multi-Functional and Biocompatible Parafilm. Micromachines (Basel), 14(3). 10.3390/mi14030656

Yuan, Q., Fang, H., Wu, X., Wu, J., Luo, X., Peng, R., Yan, S. (2023). Self-Adhesive, Biocompatible, Wearable Microfluidics with Erasable Liquid Metal Plasmonic Hotspots for Glucose Detection in Sweat. ACS Appl Mater Interfaces. 10.1021/acsami.3c11746

Zırh, S., Erol, S., Zırh, E. B., Sokmensuer, L. K., Bozdag, G., & Muftuoglu, S. F. (2021). A new isolation and culture method for granulosa cells. Cell Tissue Bank, 22(4), 719–726. 10.1007/s10561-021-09929-5

